# Kernel multitask regression for toxicogenetics

**DOI:** 10.1101/171298

**Authors:** Elsa Bernard, Yunlong Jiao, Erwan Scornet, Veronique Stoven, Thomas Walter, Jean-Philippe Vert

## Abstract

The development of high-throughput *in vitro* assays to study quantitatively the toxicity of chemical compounds on genetically characterized human-derived cell lines paves the way to *predictive toxicogenetics*, where one would be able to predict the toxicity of any particular compound on any particular individual. In this paper we present a machine learning-based approach for that purpose, kernel multitask regression (KMR), which combines chemical characterizations of molecular compounds with genetic and transcriptomic characterizations of cell lines to predict the toxicity of a given compound on a given cell line. We demonstrate the relevance of the method on the recent DREAM8 Toxicogenetics challenge, where it ranked among the best state-of-the-art models, and discuss the importance of choosing good descriptors for cell lines and chemicals.

## 1 Introduction

The toxicity of drugs or other chemicals varies between individuals. Understanding the genetic basis of this variability and being able to identify individuals prone to adverse health effects will play an increasingly important role with the development of targeted therapies in precision medicine, and could contribute to the identification of individuals at risk under specific exposure and the definition of appropriate regulatory limits for environmental health protection. *Toxicogenetics* is the name of the field that aims at understanding the genetic basis for individual differences in response to potential toxicants [21, 18, 11]. It builds upon the fast progress in our ability to genotype individuals and more generally to measure a multitude of potential biomarkers to characterize individuals, such as mutations or gene expression, together with the fast development of techniques for high-throughput screening of collections of molecular compounds on diverse human-derived cell lines.

Recent efforts to systematically characterize the molecular portraits of individuals and to assess their response to various chemicals have started to generate rich collections of data [1], paving the way to systematic analysis in order to decipher the molecular basis for response variability and to develop predictive models of individual response [3]. For example, a consortium of teams from the University of North Carolina (UNC), the National Institutes of Environmental Health Sciences (NIEHS), and the National Center for Advancing Translational Sciences (NCATS) have recently generated a large population-scale toxicity screen, testing 156 drugs and environmental chemicals on 884 cell lines derived from individuals from different geographical origins with well-characterized genotype and transcriptome [1]. These data were recently released as part of a challenge within the Dialogue on Reverse Engineering Assessment and Methods (DREAM) framework [15, 16, 4], whose goal was to derive a model to predict the response of new individuals to the various chemicals [5].

We were among the 213 participants to that toxicogenetics challenge, and describe in this paper the machine learning-based method that we developed for that purpose. It implements a kernel multitask regression (KMR), which provides a principled way to integrate heterogeneous data such as genotype and transcriptome, and to leverage information across the different chemicals tested, in a computationally efficient framework. It reached state-of-the-art performance, ranking second among 99 submissions in the first subchallenge aiming at predicting the cytotoxicity of 106 compounds across 264 cell lines that were not available during the challenge. The method is freely and publicly available^1^ as an `R` package.

## 2 Method

In this section we present the KMR predictive model we developed, based on kernel bilinear regression. We start by a brief review of positive definite kernels in the context of multitask learning, before introducing the regression model itself.

### 2.1 Kernels and multitask learning

Let 𝓧 and 𝓨 denote abstract spaces to represent, respectively, cell lines and chemicals. For example, if each cell line is characterized by a measure of *d* genetic markers, then we may take 𝓧 = ℝ^*d*^ to represent each cell line as a vector of markers, but to keep generality we simply assume that 𝓧 and 𝓨 are endowed with positive definite (p.d.) kernels, respectively *K_X_* and *K_Y_*. Remember that a p.d. kernel is a function of two arguments that generalizes the inner product, in the sense that it can be written as an inner product *K_X_*(*x*,*x′*) = Φ_*X*_(*x*)^⊤^Φ_*X*_(*y*) after mapping the data to a Hilbert space through a mapping Φ_*X*_: 𝓧 → 𝓗 [23, 27]. For example, when the data are vectors (𝓧 = ℝ^*d*^), popular p.d. kernels include the *linear kernel K*(*x*,*x′*) = *x*^⊤^*x′* and the *Gaussian Radial Basis Function (RBF) kernel with bandwidth σ: K*(*x*,*x′*) = exp (−∥*x* − *x′*∥^2^/(2*σ*^2^)). When data are graphs, such as when we represent a molecular compound by its 2D planar representation, a possible kernel is based on the comparison of the walks in the graphs [10, 7, 14]. Another interesting class of kernels is the set of *multitask* kernels, useful when we want to fit different models by sharing information across the tasks [6]. More precisely, suppose we want to fit a model of the form *f*(*x*,*y*) where *x* is a cell line and *y* is a chemical, considered as a task in the sense that we want to fit a model specific to each chemical to predict its toxicity across cell lines. Then a multitask kernel is simply a product kernel between (cell, chemical) pairs of the form *K* ((*x*,*y*), (*x′*,*y′*)) = *K_X_*(*x*, *x′*)*K_Y_*(*y*,*y′*) where *K_X_* and *K_Y_* are respectively kernels between cells and between chemicals. The standard multitask kernel of [6] interpolates between the Dirac kernel *K*(*y*,*y′*) = **1**(*y* = *y′*) and the constant kernel *K*(*y*,*y′*) = 1 by the formula *K_α_*(*y*,*y′*) = *α***1**(*y* = *y′*) + 1 − *α*, for *α* ∈ [0,1]. For *α* = 0, we get back to the constant kernel and the multitask kernel assumes in that case that all tasks are the same, and learns a single model for all tasks by pulling all data together. When *α* = 1, we get back to the Dirac kernel in which case all tasks are considered independent from each other, and a model is fit for each task without any sharing of data between the tasks. When 0 < *α* < 1, a specific model is learned for each task, but information is shared across task. When prior information is available regarding the similarities between tasks, a more general kernel between tasks can be used to share more or less information between tasks.

### 2.2 KMR

Given a set of *n* cell lines *x*_1_,…,*x_n_* ∈ 𝓧 and *p* chemicals *y*_1_,…,*y_p_* ∈ 𝓨, we assume that a quantitative measure of toxicity response *Z*_*i*,*j*_ ∈ ℝ has been measured when cell line *x_i_* is exposed to chemical *y_j_*, for *i* = 1,…,*n* and *j* = 1,…,*p*. Our goal is to estimate, from this data, a function *h*: 𝓧 × 𝓨 → ℝ to predict the response *h*(*x*, *y*) if a cell line *x* is exposed to a chemical y.

We propose to model the response with a simple bilinear regression model of the form:

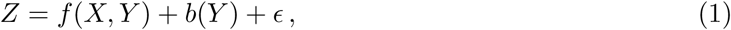

where *f* is a bilinear function, *b* is a chemical-specific bias term and *ϵ* is some Gaussian noise. We add the chemical-specific bias term to adjust for the large differences in absolute toxicity response values between chemicals, while the bilinear term *f* (*X*, *Y*) can capture some patterns of variations between cell lines shared by different chemicals. We only focus on the problem of predicting the action of known and tested chemicals on new cell lines, meaning that we do not try to estimate *b*(*Y*) on new cell lines.

If *x* and *y* are finite dimensional vectors, then the bilinear term *f* (*x*, *y*) has the simple form *x*^⊤^*My* for some matrix *M*, with Frobenius norm ∥*M*∥^2^ = *Tr*(*M*^⊤^*M*). The natural generalization of this bilinear model to possibly infinite-dimensional spaces 𝓧 and 𝓨 is to consider a function *f* in the product reproducing kernel Hilbert space 𝓗 associated to the product kernel *K_X_K_Y_*, with Hilbert-Schmitt norm ∥*f*∥^2^. To estimate model (1), we solve a standard ridge regression problem:

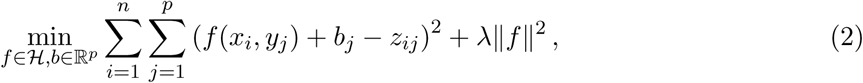

where λ is a regularization parameter to be optimized, and call the resulting method *Kernel Multitask Regression* (KMR). As shown in the next theorem, (2) has an analytical solution. Note that 1_*n*_ refers to the *n*-dimensional vector of ones, *Diag*(*u*) for a vector *u* ∈ ℝ^*n*^ refers to the *n* × *n* diagonal matrix whose diagonal is *u*, and *A* ∘ *B* for two matrices of the same size refers to their Hadamard (or entrywise) product.

#### Theorem 1.

*Let Z* ∈ ℝ^*n*×*p*^ *be the response matrix*, *and K_X_* ∈ ℝ^*n*×*n*^ *and K_Y_* ∈ ℝ^*p*×*p*^ *be the kernel Gram matrices of the n cell lines and p chemicals*, *with respective eigenvalue decompositions K_X_* =
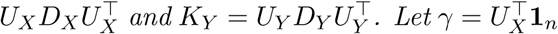
*and S* ∈ ℝ^*n*×*p*^ *be defined by*
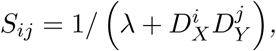
*where*
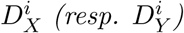
*denotes the i*-*th diagonal term of D_X_ (resp. D_Y_) Then the solution* (*f*^∗^,*b*^∗^) *of (2) is given by*

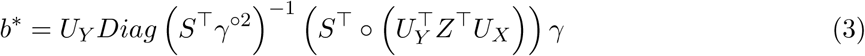

*and*

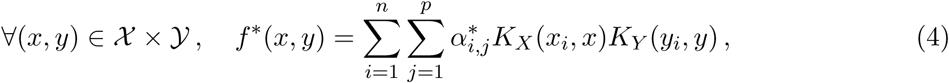

*where*

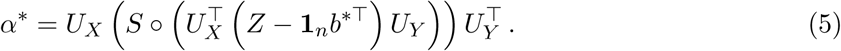

The most computationally expensive part to compute *b*^∗^ and *α*^∗^ from (3) and (5) is the eigenvalue decomposition of *K_X_* and *K_y_*, and we only need to manipulate matrices of size smaller than *n* × *n* or *p* × *p.* The computational complexity of KMR is therefore *O*(*max*(*n*,*p*)^3^), and the memory requirement *O*(*max*(*n*,*p*)^2^).

#### Proof.

By the representer theorem [22], we know that there exists a matrix *α* ∈ ℝ^*n*×*p*^ such that the solution *f* ∈ 𝓗 of (2) can be expanded as:

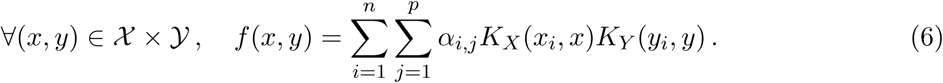

Plugging back (4) into (2) and using the fact that ∥*f*∥^2^ = *Tr*(*α*^⊤^*K_X_αK_y_*) leads to the problem:

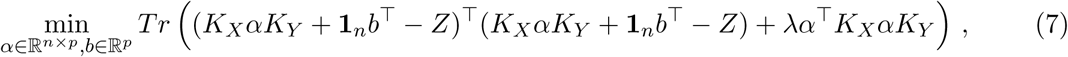

where *Tr*(*M*) is the trace of the square matrix *M*. This is a convex, quadratic program in *α* and *b*, so we can solve it by setting its gradient to zero. The gradient in *α* and *b* are respectively:

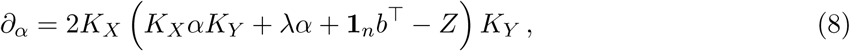

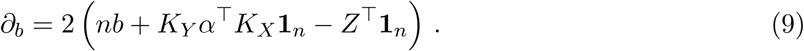

The gradient in *α* (8) is null if and only if

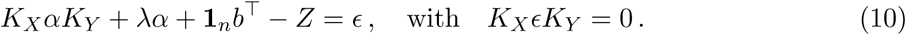

Note that although different *α* may satisfy (10) (with different *ϵ*), they all define the same function *f* through (6), since the original problem (2) is strictly convex in *f* and has therefore a unique solution. We can therefore only focus on the solution corresponding to *ϵ* = 0 in (10), leading to

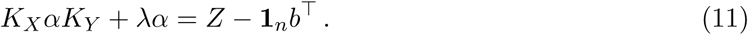

Multiplying on the left by
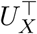
and on the right by *U_y_* leads to

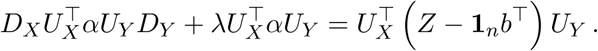

The (*i*, *j*)-th entry of the l.h.s. matrix is
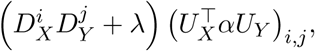
so by denoting
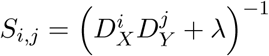
as in Theorem 1 we get:

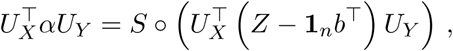

leading to

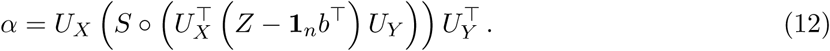

The gradient in *b* (9) is null if and only if

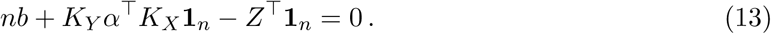

However, (11) leads to

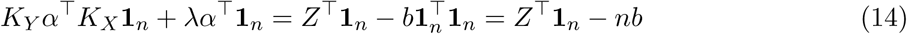

which combined with (13) gives *α*^⊤^ **1**_*n*_ = 0. Applying this condition to (12) gives

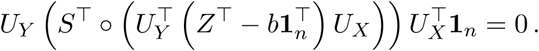

Multiplying on the left by
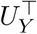
and using the notation
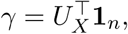
this is equivalent to

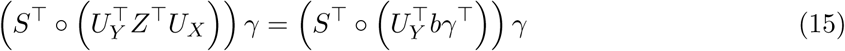

Now, if we denote
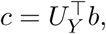
then we see that the *i*-th entry of the vector (*S*^⊤^ ∘ (*cγ*^⊤^)) *γ* is:

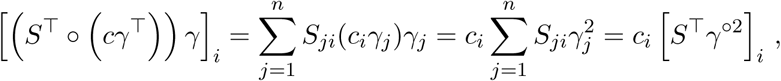

meaning

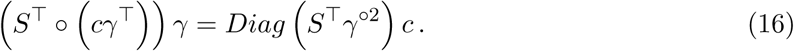

Plugging (16) into (15) we get

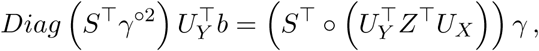

which finally gives

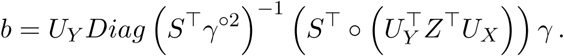

□

### 2.3 Related work

Although the explicit derivation of how to solve (2) efficiently is new to our knowledge, KMR is a natural extension of standard ridge regression to the multitask setting. Indeed, suppose first that there is a single chemical (*p* = 1) and each cell lines is represented by a *q*-dimensional vector (𝓧 = ℝ^*q*^). In that situation, if we consider the linear kernel *K*(*x*,*x′*) = *x*^⊤^*x′*, then the functions in the space 𝓗 are linear functions of the form *f* (*x*) = *w*^⊤^*x* for *w* ∈ ℝ^*q*^ and the KMR model (2) boils down to the standard ridge regression problem:

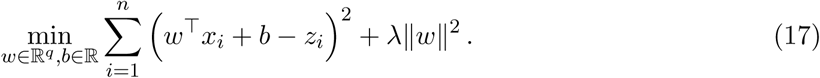

When a nonlinear kernel is used, this turns into a kernel ridge regression problem, which is a standard way to solve nonlinear regression problems [28, 23] and can also be interpreted as a mean posterior estimate of a Gaussian process [20], in which case the kernel is interpreted as a covariance function. Kernel ridge regression is also similar to support vector regression (SVR), which is obtained by replacing the squared error loss in (17) by another, so-called ∈-insensitive loss function [26], while the SVM method corresponds to another loss function called hinge loss, specifically adapted to the classification setting when the response is binary and not continuous. When several chemicals are considered (*p* > 1), KMR is equivalent to the multitask SVM formulation of [6] adapted to the regression setting by replacing the hinge loss by the squared error loss, and with a task-specific offset term. In particular, when the kernel between chemicals is the Dirac kernel, then KMR just amounts to solving *p* independent kernel ridge regression problems, one for each chemical, while for more general kernels the regression functions for each chemicals are estimated jointly. There exists a vast literature on multitask learning [2], and numerous algorithms have been proposed. While working with a product kernel as we do is standard with kernel methods, a parallel line of thoughts exists in the Gaussian process literature where hierarchical Bayesian models are used [31]. In particular, such a Bayesian formulation combining in addition an optimization of how to combine different kernels, called Bayesian multitask multiple kernel learning (MKL) was successfully used in a biological context recently [4]. Both KMR and the Bayesian multitask MKL share in common the ideas of kernelized regression, which allows multiview learning, and of multitask learning through a kernel (or covariance) between chemicals. KMR results however in a simpler and faster algorithm as the optimization problem (2) can be solved analytically in one step, while the Bayesian multitask MKL is intractable and solved approximately by a variational approximation [8].

## 3 Data

We test KMR on the data of the DREAM 8 Toxicogenetics challenge^2^, a joint crowdsourcing initiative of Sage Bionetworks, DREAM and scientists at UNC, NIEHS and NCATS to assess the possibility of developing predictive models in toxicogenetics. The effect of 156 chemical compounds was tested on 884 lymphoblastoid cell lines derived from participants in the 1000 Genomes Project and representing 9 distinct geographic subpopulations. The toxicity response was measured in terms of EC10, *i.e.*, the concentration in chemical compound at which the intracellular ATP content is decreased by 10 percent. During the challenge, participants had access to a subset of these data, corresponding to 106 chemicals tested on 620 cell lines. The goal of the challenge we focus on was to predict the effect of these 106 chemicals on the 264 cell lines that were not given to the participants. For 15 among the 106 chemicals tested, no effect on cytotoxicity was observed across the entire population [5]. To avoid the introduction of noise in the ranking, these compounds were not considered to score the predictions during the challenge, and we also removed them from our experiments below. As all training and test data were released after the challenge was over, we combine them together to obtain a set of 91 chemicals tested on 884 cell lines, the largest toxicogenetics dataset available so far.

For each cell line we have access to three covariates (population, batch and sex), to DNA variation profiles (approximately 1.3 million single nucleotide polymorphisms or SNPs), and to gene expression levels by RNA sequencing for a subset of the cell lines (46 256 transcripts). Each chemical also comes with a set of structural attributes obtained by standard chemoinformatics methodologies, including 160 descriptors based on the Chemistry Development Kit (CDK) and 9272 descriptors based on the Simplex Representation of Molecular Structure (SIRMS). In addition, we compute for each chemical 881 binary descriptors encoding the presence or absence of 881 substructures defined in the PubChem database [12], and 1554 descriptors describing the ability of the chemicals to interact with 1554 human proteins known to be potential targets for drugs and xenobiotics [29].

To use these various, heterogeneous data as inputs to KMR, we transform them into positive definite kernels for cell lines and chemicals, as follows.

- For each vector representation (vector of RNA-seq expression after log transformation, and vector of SNP for cell lines; vector of CDK descriptors, vector of SIRMS descriptors, and vector of predicted targets for chemicals), we compute 10 Gaussian RBF kernels of the form
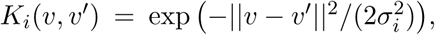
for *i* = 1,…, 10, where *v* and *v′* are two such vectors scaled such that the maximum distance ∥*v* − *v′*∥ between all pairs of cell lines (or chemicals) in the training set is equal to 1, and the bandwidth
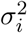
takes a value in the grid {1/20, 1/10, 1/5, 2/5, 3/5, 4/5, 1, 2, 4, 8}. We additionally compute an 11-th kernel as the mean of the 10 Gaussian RBF kernels, namely,
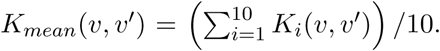
This results in a total of 11 kernels for each vector representation, giving 22 cell line kernels (denoted `Krnaseq∗` and `Ksnp∗` for respectively the kernels based on RNA-seq and SNP data) and 33 chemical kernels (denoted respectively `Kcdk∗`, `Ksirms∗` and `Kpredtarget∗` for the kernels based on CDK, SIRMS and predicted target descriptors). In each kernel name, the `∗` character refers to the string `Rbf1-Rbf10` to represent a Gaussian RBF kernel with one of the 10 bandwidths, or **Mean** to represent the mean of the 10 Gaussian RBF kernels. For example, `KrnaseqRbf3` refers to the Gaussian RBF kernel on cell lines based on RNA-seq data with bandwidth *σ*^2^ = 1/5, while `KcdkMean` refers to the kernel for chemicals obtained by averaging the 10 Gaussian RBF kernels built from CDK descriptors.
- From the 3 covariates for cell lines (population, sex and batch), we build 4 kernels: one linear kernel for each covariate, and one linear kernel for the vector of covariates. This amounts to fitting a linear model based on a single covariates, or on all covariates together. We denote these kernels respectively by `KcovariatesPopulation`, `KcovariatesSex`, and `KcovariatesBatch` for the kernels based on each individual covariate, and `Kcovariates` for the kernel based on all covariates together.
- For chemicals, we include two kernels based on detecting substructures in their 2D representations: a marginalized graph kernel [10] as implemented in the `ChemCPP` package with defaults parameters^3^, and a linear kernel obtained from the PubChem fingerprint which characterizes each chemical compound by a binary vector indicating the presence or absence of 881 substructures [12]. We denote these kernels respectively `Kchemcpp` and `Ksubstructure`.
- We also consider a series of 11 plain multitask kernels for chemicals, of the form *K_α_*(*y*,*y′*) = *α* + (1 − *α*)**1**(*y* = *y′*), for *α* ∈ {0, 0.1, 0.2, 0.3, 0.4, 0.5, 0.6, 0.7, 0.8, 0.9, 1}, and denote them by `Kmultitask∗` where `∗` is a number from 1 to 11 to indicated the value of *α* in the grid. For example, `Kmultitask1` refers to *α* = 0 hence the Dirac kernel *K*_0_(*y*,*y′*) = **1**(*y* = *y′*); this amounts to considering all chemicals independently from each other, and fitting a model for each chemical without sharing information between chemicals. The other extreme is `Kmultitask11`, corresponding to *α* = 1, hence the constant kernel *K*_1_(*y*, *y′*) = 1; this amounts to pooling together all chemicals and fitting a single model for toxicity for all compounds. In between, `Kmultitask6` is the standard multitask kernel *K*_1/2_(*y*,*y′*) = (1 + **1**(*y* = *y′*)) /2 of [6] which allows to share information across chemicals, while still fitting specific models for each chemical.
- We also define an empirical kernel between chemicals, denoted `Kempirical` and equal to the correlation between the toxicity profiles of any two chemicals on the training set. Like the multitask kernels, the empirical kernel does not use any chemical descriptors, but instead promotes the sharing of information between chemicals that appear to have similar profiles on the training set.
- Finally, for both cell lines and chemicals, we consider an integrated kernel defined as the sum of all other kernels. Summing kernel is a simple and popular way to combine different heterogeneous descriptors [19, 30]. We denote it by `Kint`.

In total, this results in 27 cell line kernels and 48 chemical kernels, as summarized in Table 1.

**Table 1:** List of kernels used in the study. See text for details about each kernel.

| Type | Data | Description | Name | Number |
| --- | --- | --- | --- | --- |
| Cell lines | SNP | Gaussian RBF kernels and their average | Ksnp* | 11 |
|  | RNA-seq | Gaussian RBF kernels and their average | Krnaseq* | 11 |
|  | Covariates | Linear kernels | Kcovariates* | 4 |
|  | All | Integrated kernel | Kint | 1 |
| Chemicals | CDK descriptors | Gaussian RBF kernels and their average | Kcdk* | 11 |
|  | SIRMS descriptors | Gaussian RBF kernels and their average | Ksirms* | 11 |
|  | Predicted targets | Gaussian RBF kernels and their average | Kpredtargets* | 11 |
|  | 2D structure | Marginalized graph kernel | Kchemcpp | 1 |
|  | 2D structure | Pubchem fingerprint | Ksubstructure | 1 |
|  | None | Multitask kernels | Kmultitask* | 11 |
|  | Toxicity | Empirical kernel | Kempirical | 1 |
|  | All | Integrated kernel | Kint | 1 |

## 4 Results

As explained in the Methods section and illustrated in Figure 1, we developed a machine learning-based method called KMR to predict the toxicity of a panel of chemicals on different cell lines. KMR predicts the toxicity of a chemical on a cell line by leveraging different pieces of information about the chemical and the cell line encoded in a unified and computationally efficient framework through p.d. kernels. In this section we assess empirically the performance of KMR on toxicity prediction in the context of the DREAM8 toxicogenetics challenge.

**Figure 1:**
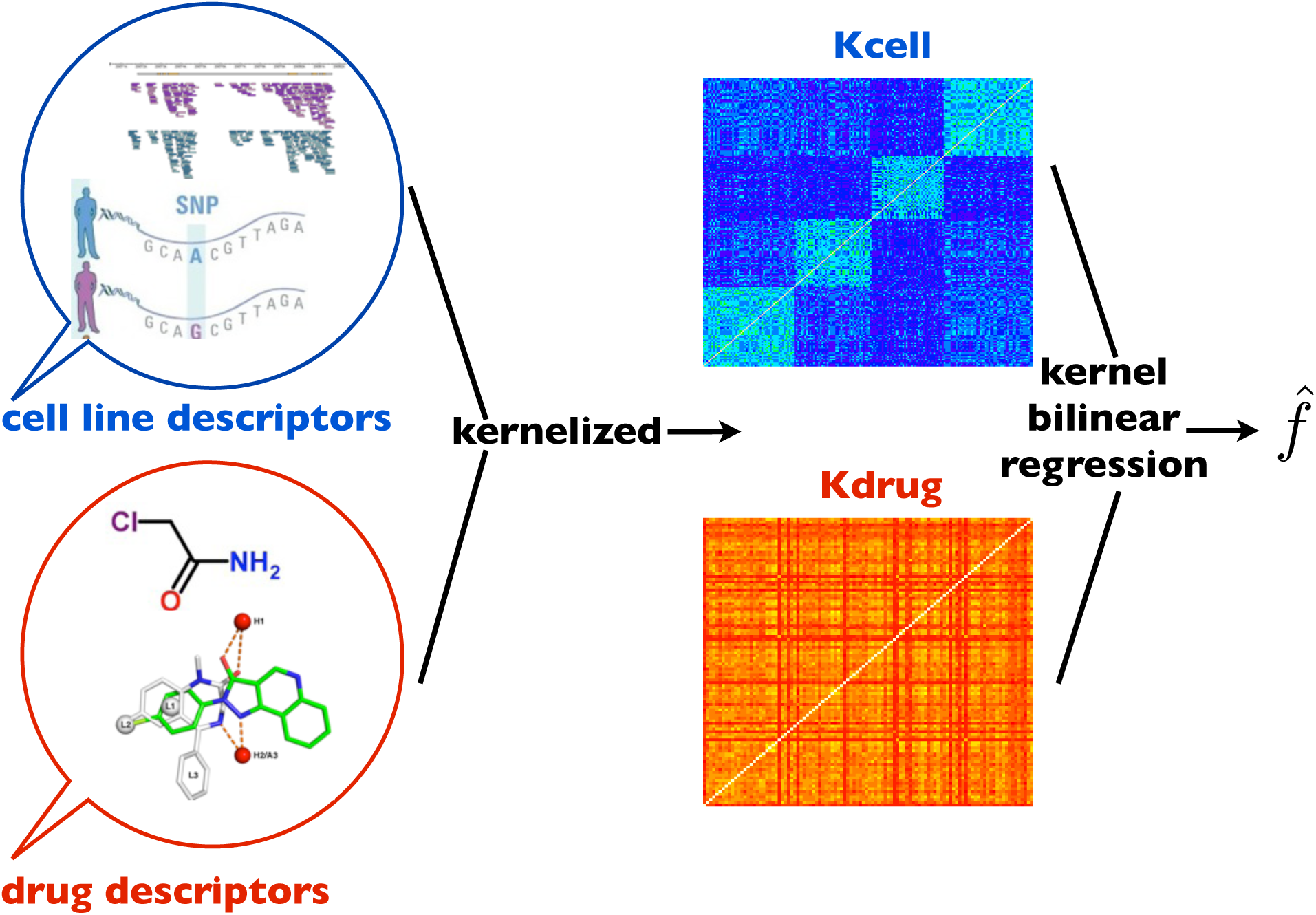
Overview of the kernel multitask regression method. Various descriptors of chemicals and cell lines are summarized into a chemical and a cell line kernel function, respectively, and a bilinear regression is performed based on the kernels used to predict toxicity of a chemical on a cell line.

### 4.1 Parameter tuning and kernel selection

Before looking at the predictive performance of KMR, let us first clarify how to tune its parameters. As seen in (2), KMR has a single parameter to tune, namely the λ regularization parameter. A large value for λ should typically be used for noisy data, or when the size of the training set is small; while a small value may be sufficient for less noisy data with a large training set. A natural way to automatically tune λ empirically is to perform cross-validation (CV) on the training set with different values of λ, and pick the value that leads to the best CV performance. Note, however, that if the toxicity profiles of chemicals have different signal-to-noise ratio, the optimal λ value may differ for different chemicals. For example, an ”easy” chemical where toxicity is easily predicted from a few genomic data may be better estimated with a small λ, while a ”difficult” one where variations in toxicity are mostly due to noise may be better predicted with a large λ. For that reason, we recommend to optimize λ for each task independently, by optimizing each task-specific performance during cross-validation. To illustrate this, Figure 2 shows the 5-fold CV performance of KMR (measured as the mean concordance index CI between the predicted and true toxicity on the CV test sets) for each chemical, as a function of λ taken over a broad grid of values {*e*^−15^, *e*^−14^,…, *e*^25^}. For this particular example we ran KMR with the integrated kernel `Kint` for cell lines, and the empirical kernel `Kempirical` for chemicals. We see that for most chemicals the performance curve is not flat, suggesting that tuning λ is important; we also see that the optimal λ value estimated for each chemical (the black circles in Figure 2) differs significantly between chemicals, suggesting that it can be better to pick different λ’s for different chemical. From a computational point of view, note that it is not more expensive to tune different λ’s for different chemicals than to tune a single λ for all chemicals; in both cases a KMR model has to be trained on each CV training set for each value of λ, the only difference being that in the first case we maximize each curve on Figure 2, while in the second case we maximize the mean of the curves.

**Figure 2:**
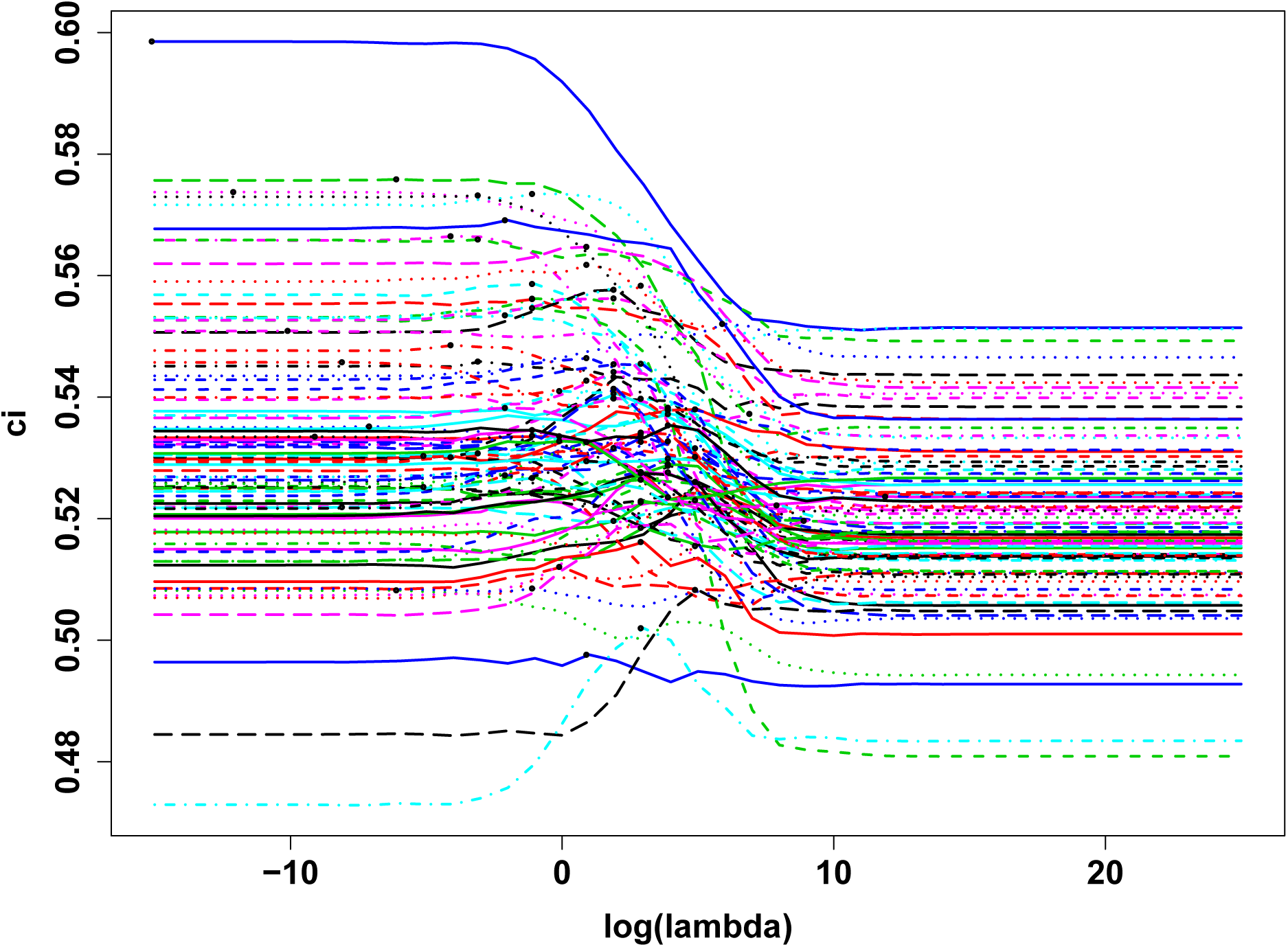
Cross-validation performance of KMR for different chemicals (each curve corresponds to one chemical) as a function of λ. The performance is measured in mean concordance index over the 5 test sets of a 5-fold cross-validation run. Black circles indicate, for each chemical, the optimal λ estimated.

The other, important hyperparameters to choose for KMR are the kernels used for cell lines and chemicals. As this is akin to the more general problem of which features to use to characterize cell lines and chemicals, and how to engineer good features to predict toxicity, we have no better advice than to rely on one’s intuition and to test different kernels empirically. When different kernels can be thought of, we note that a simple way to combine them is to sum them together (what we refer to as the integrated kernel `Kint`), providing a simple default solution if one wants to pick a single kernel to summarize heterogeneous data such as genotype, gene expression and covariates for cell lines. As for chemicals, which correspond to tasks in the challenge, we note that some kernels can depend on descriptions about the chemicals (such as their 2D structure), while other kernels such as the multitask kernel `Kmultitask∗` and the empirical kernel `Kempirical` use no information about the chemicals. Using chemical information may be relevant if one believes that a particular chemical feature is important for toxicity, and if one wants to be able to make toxicity predictions for chemicals not seen in the training set; however, if we just want to make toxicity predictions for compounds of the training set, `Kempirical` and `Kmultitask∗` may be good default kernel. Based on these considerations, we provide below an empirical comparison of the different kernels we investigated during the challenge, and then discuss the challenge results where we submitted a KMR model based on the integrated kernel **Kint** for cell lines and the empirical kernel **Kempirical** for chemicals.

### 4.2 Performance on the DREAM8 toxicogenetics dataset

We now assess the performance of KMR to predict the toxicity of known chemicals on new cell lines on the data of the DREAM8 Toxicogenetics challenge. As explained above, we collected from the challenge 80,444 EC10 toxicity measurements of 91 chemicals on 884 cell lines. To mimic the setting of the challenge on this dataset, we assess the performance of different methods by 5-fold cross-validation over the cell lines, repeated 10 times, meaning that we split 50 times the cell lines into a training set (80% of the cell lines) used to train a model, and a test set (20% of the cell lines) used to assess the performance. Following the method used in the challenge, we assess the performance of prediction in terms of concordance index (CI) on the test set per chemical, averaged over the chemicals. A random prediction leads to a CI of 0.5, while a CI of 1 means that we have perfectly ranked the cell lines in terms of response to the chemicals.

As explained above, we explored different ways to represent the cell lines and chemicals through different p.d. kernels, and the kernel multitask model can take as input any of the 27 cell line kernels in combination with any of the 48 chemical kernels, resulting in 27 × 48 = 1, 296 combinations. Note that only 337 cell lines out of 884 had RNA-seq information; for cell lines missing RNA-seq information, we replaced any RNA-seq-based kernel by the Dirac kernel. Figure 3 summarizes the performance reached by KMR with the different combinations of kernels. A first, disappointing observation is that the overall performance barely reaches *CI* = 0.54, which is significantly better than random but small in absolute value. However, this is coherent with the global *a posteriori* analysis of the challenge that confirmed that the task was difficult and that the performance of all models remains limited [5].

**Figure 3:**
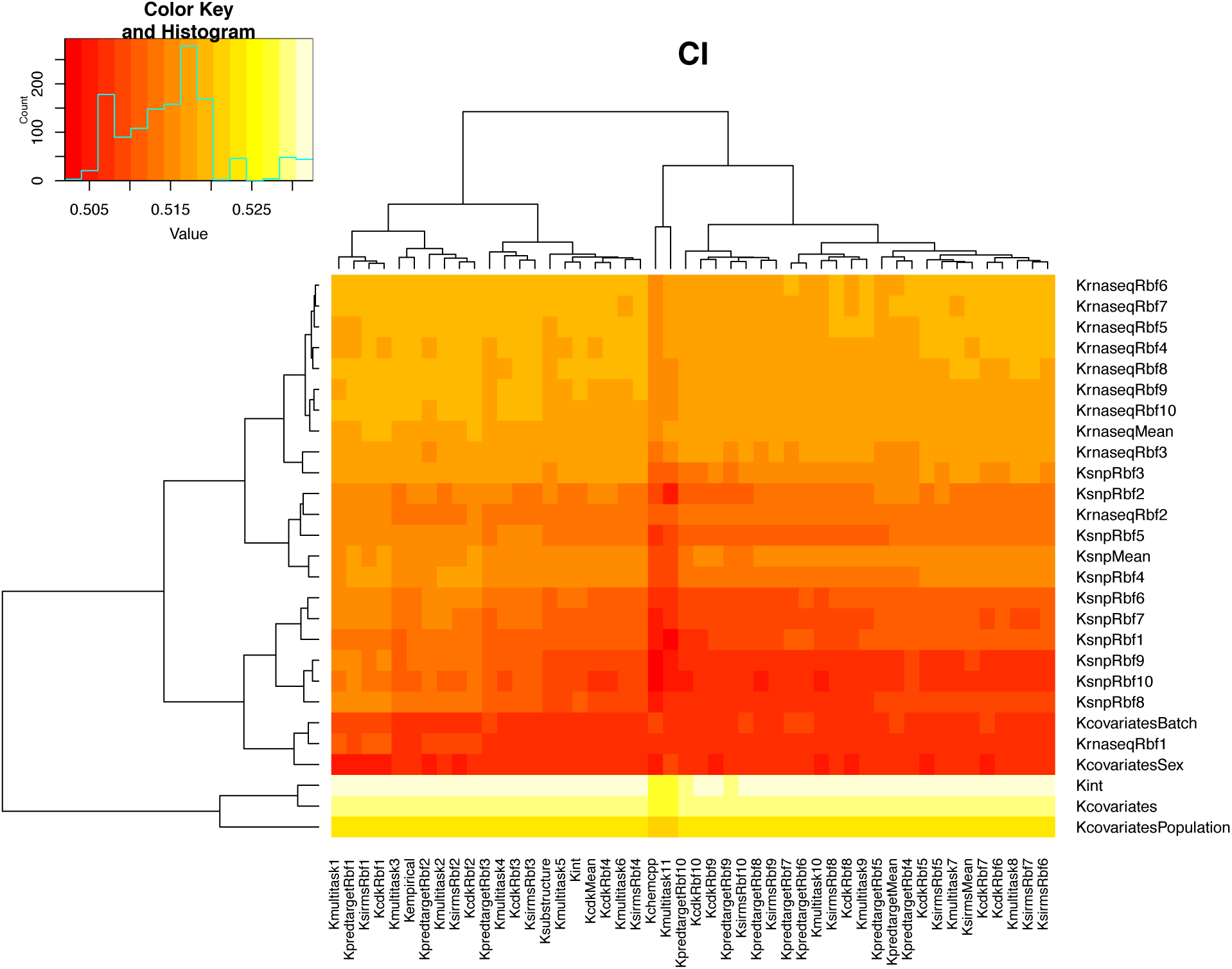
Mean CI for each combination of cell line kernel (vertical axis) and chemical kernel (horizontal axis), by cross-validation over the full set of 884 cell lines.

This being said, we observe clear differences between the performances of different kernels, particularly for cell line kernels. To highlight the impact of the different kernels, we show on Figures 4 and 5 the mean performance reached by each cell line kernel, respectively each chemical kernel. Interestingly, the best cell line kernel is `Kint`, the integrated kernel, with mean CI above 0.53. The fact that it outperforms each kernel based on a single data source suggests that there is relevant information about toxicity spread over different types of cell line data, and confirms that the integrated kernel is a simple and powerful approach to integrate heterogeneous information for prediction purpose. Regarding cell line kernels based on a single type of data, the best results are obtained with the covariates, particularly the population information; followed by gene expression, as long as the bandwidth of the Gaussian kernel is not too small. SNP-based kernels have slightly lower performance, and are more sensitive to the bandwidth of the kernel; interestingly, the ”mean” kernel for both RNA-seq and SNP, which averages the Gaussian kernels with different bandwidth, perform in both cases almost as well as the kernel with the best bandwidth, suggesting that the ”mean” kernel is a good alternative to optimizing the bandwidth. On the chemical side, we see much less influence of kernels. The kernels that give the best performances, with a small margin, tend to be Gaussian kernels with small bandwidths, or the Dirac kernel (`Kmultitaskl`) which corresponds to a Gaussian kernel with zero bandwidth. This suggests that on this dataset, there is little benefit to share information between chemicals. For example, the performance of the plain multitask kernels `Kmultitask∗` keeps degrading slightly as more information is shared between tasks. The only kernel near the top of the list that differs a lot from the Dirac kernel is the empirical kernel, which could be capturing some interesting signals between chemicals with similar toxicity profiles on the training set.

**Figure 4:**
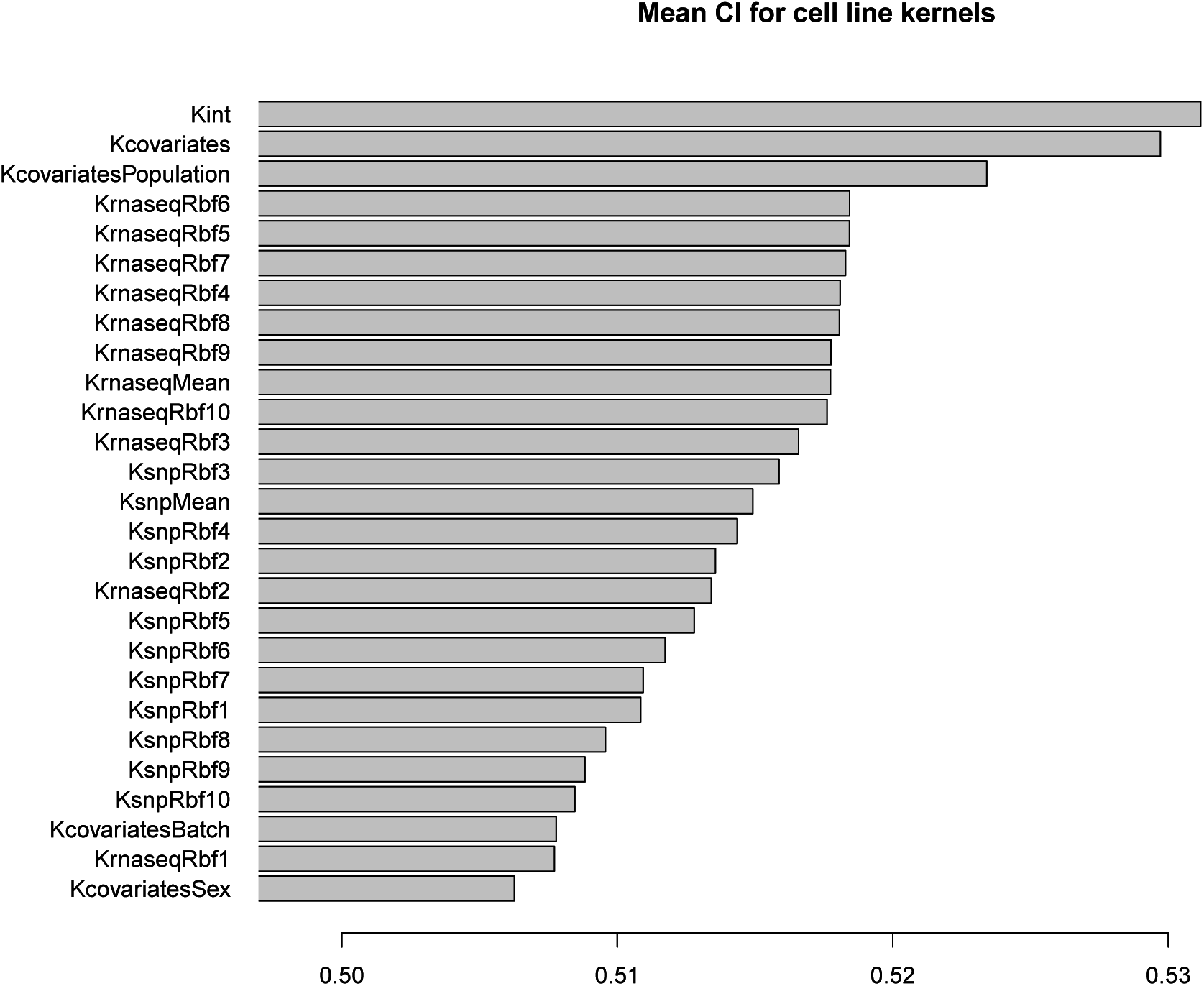
Average CI reached by each cell line kernel, on the 884 cell lines.

**Figure 5:**
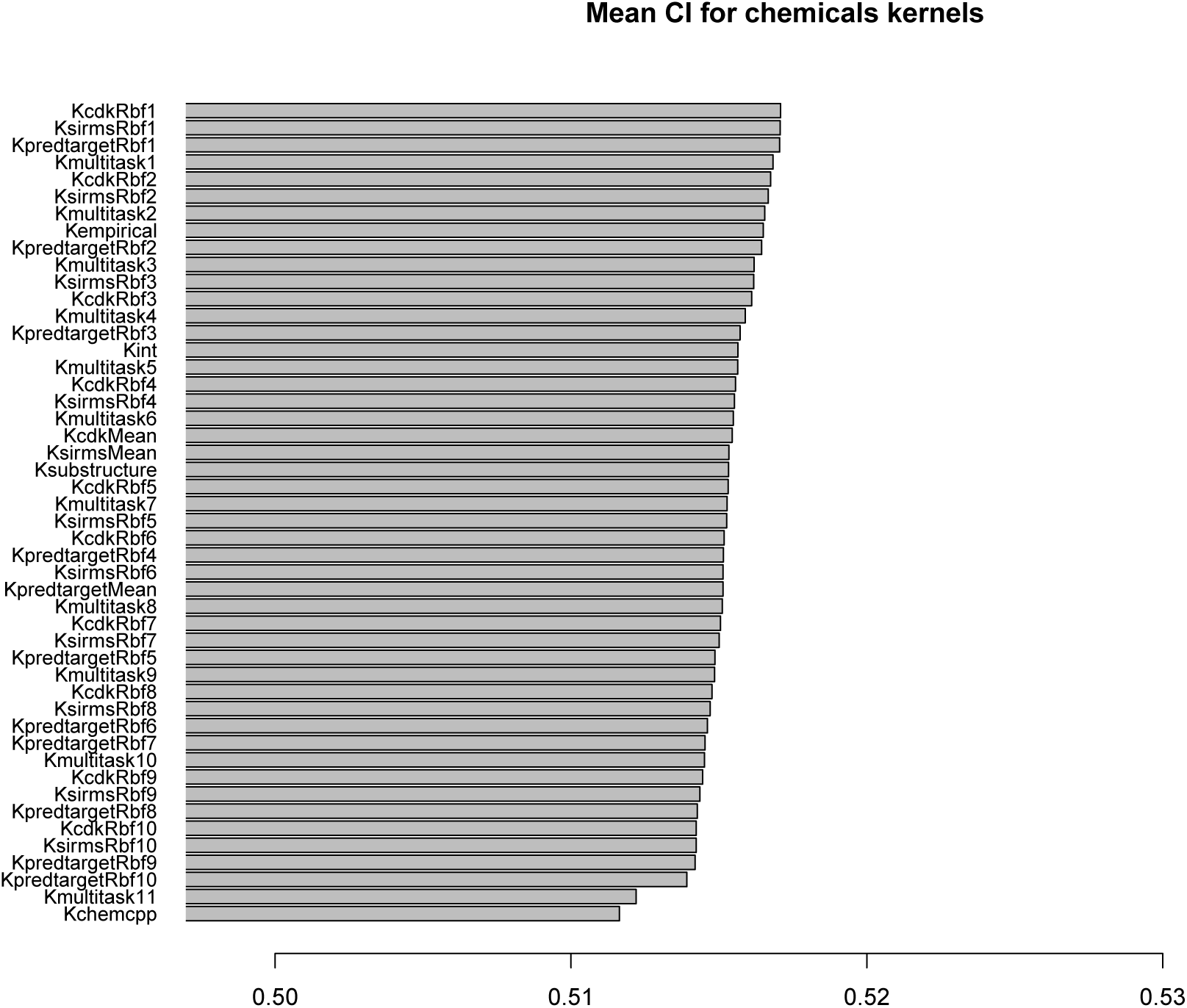
Average CI reached by each chemical kernel, on the 884 cell lines.

Since the RNA-seq data was only available on 337 cell lines, we also analyze the behavior of KMR by cross-validation on the subset of cell lines with RNA-seq data (Figure 6). In spite of the reduced set of the training set, we notice no decrease in performance compared to the results on the full set of cell lines (Figure 3), if not a slight increase. The relative performance of the different kernels is overall conserved compared to the experiment on all cell lines, except that, unsurprisingly, the performance of kernels based on RNA-seq information increases and, for example, now outperforms the performance of the covariate kernel. However, the integrated kernel remains the best overall.

**Figure 6:**
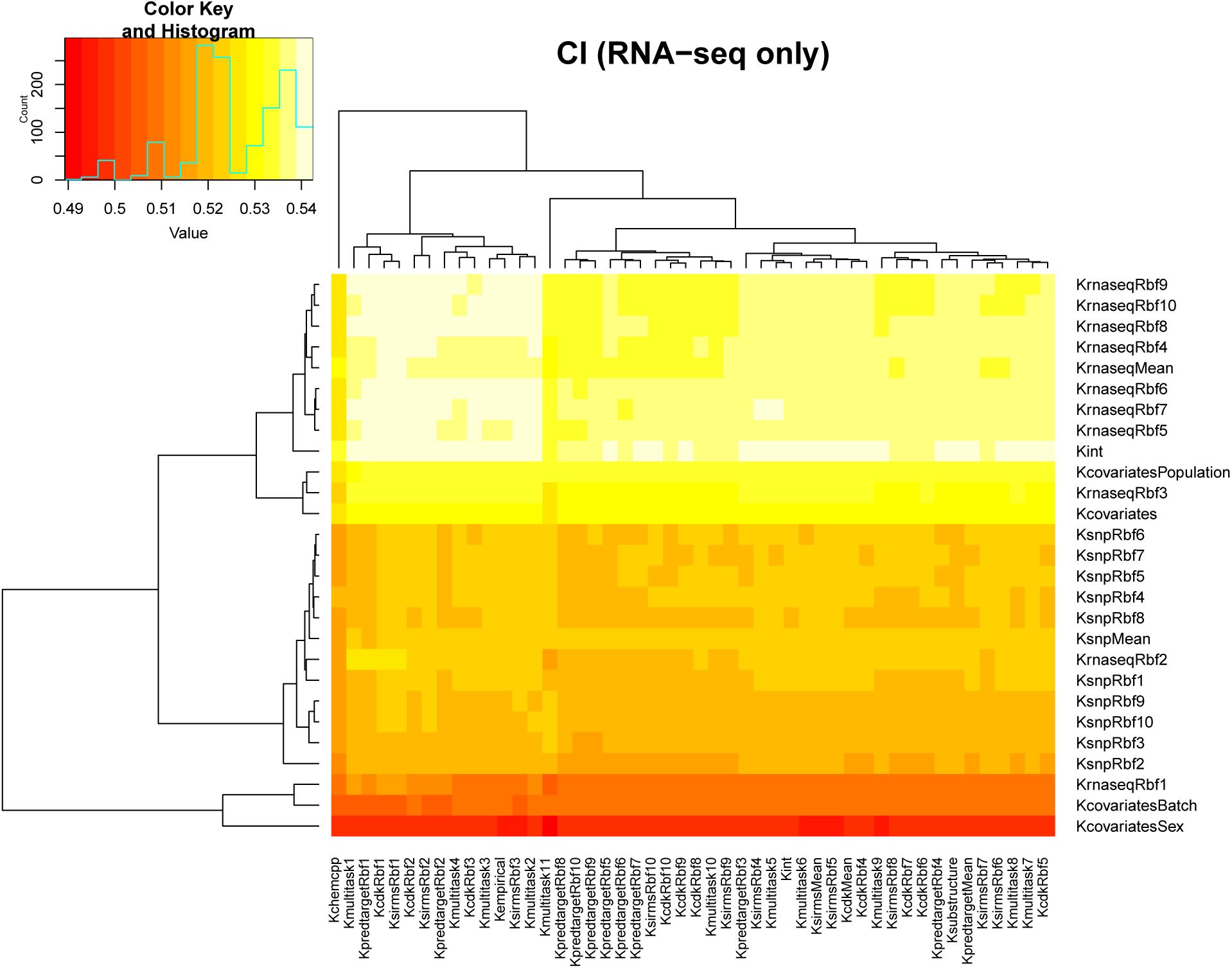
Mean CI for each combination of cell line kernel (vertical axis) and chemical kernel (horizontal axis), by cross-validation over the set of 337 cell lines with RNA-seq available.

To assess the performance of KMR compared to other, standard methods for regression, we compare it to regression with elastic net, lasso and random forests [9]. Since these methods have no obvious multitask version, we run them in a single task mode, fitting a model on each chemical independently. For all methods, we give as input the `Kint` matrix to represent cell lines; for KMR, we additionally use `Kempirical` to share information among chemicals. We note that not using a multitask formulation is probably not a strong disadvantage on this dataset, as we showed that the Dirac kernel has a good performance with KMR, and as the winner of the challenge trained a RF model for each chemical separately, with specifically handcrafted features. Here our goal is just to get a sense of how different methods behave if given the same input data. Figure 7 summarizes the distribution of CI values over the 91 chemicals for the different methods. While elastic net, lasso and RF regression are not significantly different, KMR significantly outperforms them (paired t-test p-value < 10^−13^). Furthermore, KMR is about 30 times faster than lasso and elastic net, and more than 100 times faster than RF trained with only 100 trees (Table 2).

**Figure 7:**
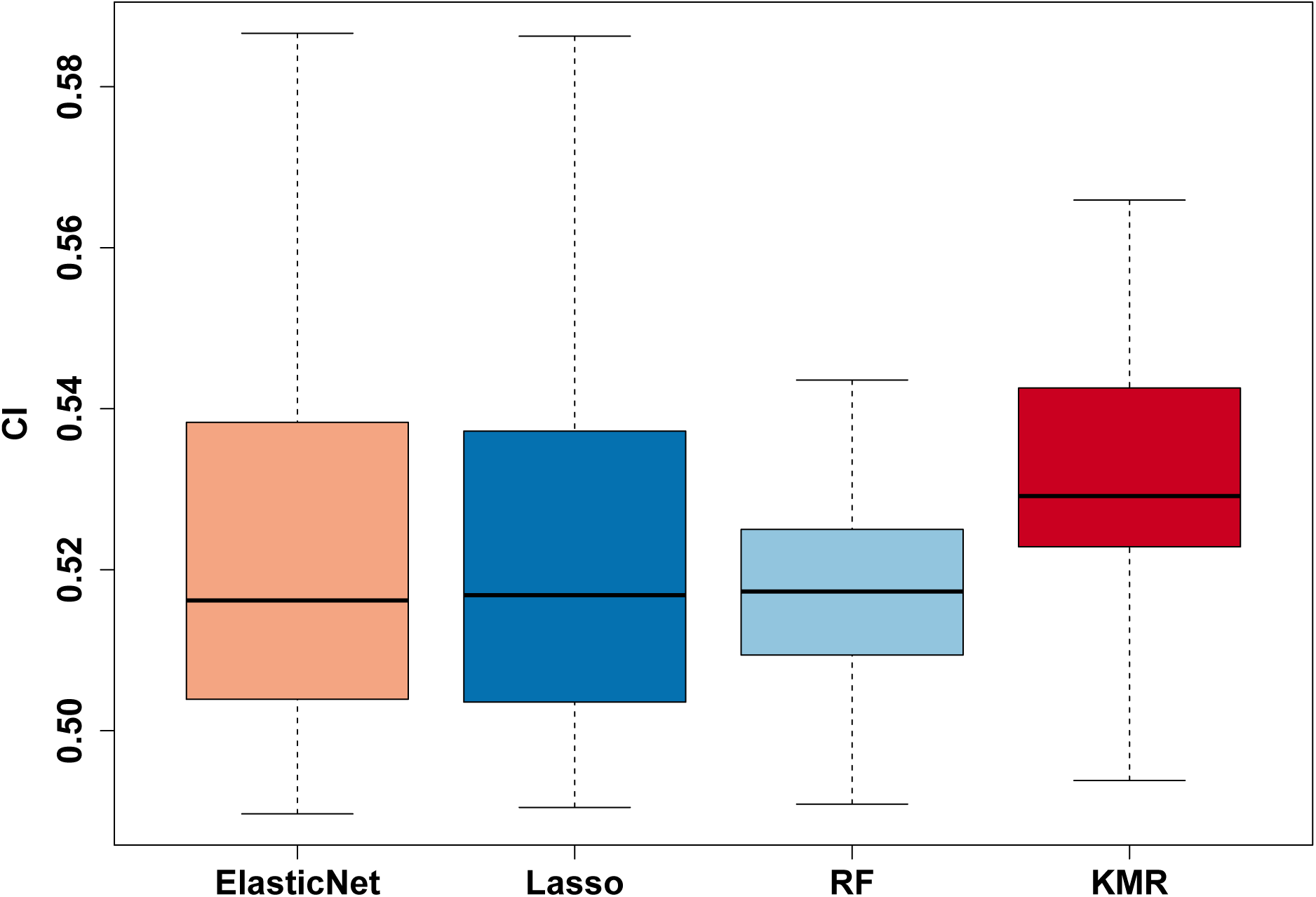
Distribution of cross-validation CI over the 91 chemicals for different methods, trained on the integrated kernel matrix to represent cell lines (and the empirical kernel between chemicals for KMR).

**Table 2:** CPU time to train a model on half of the full dataset, including parameter optimization by cross-validation, and to predict the toxicity on the second half. All methods were run from `R`, using the `glmnet` package for elastic net and lasso, the `randomForest` package for random forests, and the `kmr` package for KMR. All methods were used with default parameters, except for the number of trees of the random forest which was reduced from the default `ntree=500` to `ntree=100` to reduce by a factor of 5 the total CPU time, which is already two orders of magnitude slower than KMR.

| Method | Time (s.) |
| --- | --- |
| Elastic net | 262 |
| Lasso | 263 |
| Random forest | 1,173 |
| KMR | 9 |

In summary, this analysis by cross-validation on DREAM toxicogenetics dataset of the different combinations of kernels shows that (i) the absolute performance remains limited but varies with the combination of kernels used, (ii) there is some predictive information in the population information, and in the RNA-seq information, (iii) the integrated kernel Kint consistently outperforms all individual cell line kernels, (iv) there is limited benefit of using a multitask learning strategy, and (v) KMR is a fast and competitive regression approach in this context compared to state-of-the-art methods.

### 4.3 Results during the challenge

We participated as a team named *CASSIS* to the DREAM Toxicogenetics challenge. During the challenge, we only had access to toxicity measurements for a training set of 620 cell lines, and each participant had to submit toxicity predictions for the remaining 264 cell lines. submissions were scored and ranked by decreasing mean CI over the 91 chemicals. We performed an analysis similar to the one described in the previous section, but restricted to the training set, in order to inform our submission. Results were overall similar, except for a strong batch effect that was present in the training set but, as shown above, is much less marked when all data are considered together. We submitted a prediction based on KMR using `Kint` for cell lines, and `Kempirical` for chemicals. The prediction ranked second out of 99 submissions in the final evaluation, outperformed only by a method based on random forests which included prior knowledge-driven selection of candidate SNP [5]. A number of different modeling strategies were used by the participants, in terms of feature engineering, data reduction, prediction algorithms and techniques for model validation. The challenge organizers assessed these strategies by surveying participants, but found no clear indication as particularly good or bad approaches, concluding that ”performance was dependent mainly on strategy for methodological application rather than on algorithmic choice” [5]. We note that our submission was the only one among the top 10 which required no feature selection and no dimension reduction, according to [5], suggesting that KMR provides a relatively simple, fast, reproducible and state-of-the-art approach.

## 5 Conclusion

We presented KMR, a new machine learning-based multitask regression model to learn simultaneously a toxicity model for different chemicals, and demonstrated through our participation and good performance at the NIEHS-NCATS-UNC DREAM Toxicogenetics challenge that it reaches state-of-the-art performance on toxicity prediction across cell lines. The model is modular in the sense that it can combine any description of the cell lines with any description of the chemicals, using the kernel trick. We investigated the relevance of several kernels based on genomic and chemical information, but more kernels could be designed and explored based on the rich literature of kernel design [24]; for example, position-specific kernels for SNP or kernels based on copy number variations could enrich the encoding of genomic information used to characterize cell lines. KMR is completely generic, implemented as a publicly available `R` package, and could be used for other multitask regression problems such as predicting the response to a panel of drugs based on the genetic and molecular characterization of cancers [4]. Extensions of the model to predict not only a single real-valued output but a more complex output, such as a vector, is an interesting future work that falls in the theory of structured-output regression with kernels [25, 17]; this could, for example, model the variations in the transcriptome of different cell lines subject to different chemicals, as experimentally assayed in the Connectivity Map project [13].

## 6 Acknowledgements

We thank an anonymous reviewer for several suggestions which considerably improved the presentation of our work. This work was supported by the European Union 7th Framework Program through the Marie Curie ITN MLPM grant No 316861, and by the European Research Council grant ERC-SMAC-280032. The authors declare no conflict of interest that might bias their work.

## Footnotes

1 https://github.com/jpvert/kmr

2 https://www.synapse.org/#!Synapse:syn1761567

3 http://chemcpp.sourceforge.net

